# Intrinsic-like blood coagulation activity in fish: Involvement of hepatocyte growth factor activator and interspecific diversity

**DOI:** 10.64898/2026.10.23.761479

**Authors:** Shintaro Matsui, Asami Yoshida

## Abstract

Most fish lack identifiable orthologs of factor XII and other components of the mammalian contact system; however, their plasma exhibits coagulation responses to negatively charged surfaces. Therefore, hepatocyte growth factor activator (HGFA), an evolutionary relative of factor XII, has been proposed as an initiator of factor XII-independent, contact activation-like coagulation in fish. To examine the distribution of this activity, we compared plasma from multiple fish species using an activated partial thromboplastin time (aPTT)-like assay. Thrombin substrate-cleaving activity was detected under aPTT-like conditions in multiple species lacking identifiable factor XII orthologs, demonstrating that factor XII-independent intrinsic pathway-like activity is broadly distributed among fish. To assess the involvement of factor XII- or HGFA-related proteases, we performed inhibitor assays. The observed inhibitor sensitivities did not consistently support HGFA as the principal initiator of the activity. We next compared the occurrence and intensity of the activity with the genomic presence and domain organization of HGFA orthologs. Neither the presence nor the magnitude of the activity closely corresponded to HGFA occurrence or structure; notably, activity was detected in bichir, in which no clear HGFA ortholog has been identified. Finally, comparative domain analysis showed that fish HGFA generally lacks several structural features associated with the surface-dependent autoactivation of mammalian factor XII. These findings indicate that factor XII-independent, contact activation-like coagulation is broadly present in fish but cannot be explained solely by HGFA. Fish may therefore possess diverse or unidentified proteolytic mechanisms that initiate coagulation under aPTT-like conditions, providing insights into the evolutionary emergence of the vertebrate contact system.

**Highlights:** Intrinsic-like coagulation activity was detected in some fish species.

The sensitivity to corn trypsin inhibitor differed among fish species.

Factor Xa bypasses the inhibitor action upstream of the coagulation cascade.

Models of hepatocyte growth factor activator and inhibitor showed binding modes.

The activator domain structure disfavors factor XII-like autoactivation.

## Introduction

In the classical cascade model, plasma coagulation is described in terms of the extrinsic and intrinsic pathways. The extrinsic pathway is initiated by the tissue factor, whereas the intrinsic pathway is initiated in vitro by the contact activation of factor XII on negatively charged surfaces and involves factors XI, IX, and VIII (Hoffman and Monroe, 2001). Endogenous polyanions, including platelet-derived polyphosphate and extracellular nucleic acids, support factor XII-dependent contact reactions in vivo (Kannemeier et al., 2007; Müller et al., 2009). Nevertheless, congenital factor XII deficiency in humans and mice causes marked prolongation of activated partial thromboplastin time (aPTT) without an overt bleeding tendency, indicating that factor XII is dispensable for normal hemostasis (Davie and Ratnoff, 1964; Macfarlane, 1964; Park and Park, 2024). Factor XII is recognized as a component of the contact system, along with factor XI, prekallikrein, and high-molecular-weight kininogen, which contribute to thrombosis, inflammation, vascular responses, and innate immunity (Long et al., 2016; Pathak et al., 2018).

Orthologs of many major mammalian coagulation factors, including fibrinogen, prothrombin, and factor X, have been identified in fish (Hanumanthaiah et al., 2002; Jiang and Doolittle, 2003; Matsui et al., 2023). Tissue factor and factor VII orthologs are also present, and the functional conservation of tissue factor-dependent coagulation has been demonstrated in zebrafish (Beeler et al., 2018; Sheehan et al., 2001). In contrast, clear factor XII orthologs are absent from most fish species, with the notable exception of lungfish (Chana-Muñoz et al., 2019; Jiang and Doolittle, 2003; Ponczek et al., 2008). Most ray-finned fishes also lack clear orthologs of factor XI, prekallikrein, and typical high-molecular-weight kininogen, although plasma kallikrein/factor XI-like genes have been reported in some cartilaginous and bony fishes (Ponczek et al., 2008; Ponczek et al. 2020; Tsutsui et al., 2023; Wong and Takei, 2013). Thus, fish generally retain the major components of the common and extrinsic pathways but lack a complete mammalian-type contact system. Despite this molecular composition, plasma from teleosts, including zebrafish, common carp, and Japanese flounder, coagulates in response to intrinsic pathway reagents containing contact activators (Dhinoja et al., 2024; Kawatsu, 1986; Matsui, 2026). This apparent discrepancy suggests that fish lacking an identifiable factor XII ortholog may possess an alternative mechanism that promotes coagulation in response to negatively charged surfaces.

Hepatocyte growth factor activator (HGFA) has been proposed as a candidate initiator of factor XII-independent contact activation-like reactions in fish. In zebrafish plasma, a 26-nucleotide fragment derived from the 3′ terminus of erythrocyte 5.8S rRNA promoted coagulation, and this activity was attenuated by morpholino-mediated knockdown of HGFA (Alharbi et al., 2021). Because factor XII has been proposed to have arisen through duplication of an ancestral HGFA gene, zebrafish HGFA was suggested to function as a factor XII-like initiator (Ponczek et al., 2008). However, mammalian pro-HGFA is activated proteolytically by thrombin and kallikrein-related peptidases, and neither surface-dependent autoactivation nor activation of a known coagulation factor zymogen has been demonstrated for HGFA (Mukai et al., 2008; Shimomura et al., 1993). Moreover, whereas factor XII functions in the contact system activation, the established function of HGFA is to activate pro-HGF and thereby regulate HGF–MET signaling involved in tissue repair and regeneration (Fukushima et al., 2018; Miyazawa et al., 1996). Whether fish HGFA can undergo surface-dependent activation and initiate coagulation in a manner analogous to factor XII remains unknown.

In the present study, plasma responses to an aPTT reagent were compared among multiple fish species, and inhibitor assays were performed to characterize the molecular pathways underlying contact activation-like coagulation. We further examined the relationship between this activity and the presence, absence, and structural characteristics of fish HGFA orthologs, focusing on domains associated with surface-dependent autoactivation of mammalian factor XII. This study aimed to clarify the molecular basis of factor XII-independent, contact activation-like coagulation in fish and to provide insight into the evolutionary emergence of the mammalian contact system.

## 2. Materials and Methods

### 2.1 Plasma preparation

Farmed Japanese flounders (*Paralichthys olivaceus*, total length 35–45 cm) and common carp (*Cyprinus carpio*, total length 60–70 cm) were obtained from aquaculture suppliers in Fukuoka City, Japan. Gray bichir (*Polypterus senegalus*, total length 30–40 cm) were purchased from a pet shop in Tokyo, Japan, and wild bullhead sharks (*Heterodontus japonicus*, total length 50–70 cm) were acquired from a breeder in Numazu City, Japan.

Blood samples were drawn from the caudal vein of each fish and anticoagulated by mixing 9.0 mL of blood with 1.0 mL of 3.2% sodium citrate (FUJIFILM Wako Pure Chemical Corporation, Osaka, Japan). To eliminate any remaining blood cells, the samples were centrifuged at 8,000 *× g* for 10 min. The resulting plasma was stored at −80°C until further analysis. All experimental procedures were approved by the Animal Ethics Committee of Kyushu Sangyo University and were conducted in compliance with institutional guidelines (Approval No. G2025-12).

### 2.2 Enzyme inhibitors and substrates

The chromogenic substrate Nα-(2-hydroxy-3,4-difluorophenylacetyl)-D-phenylalanyl-L-arginine p-nitroanilide (HD-FPR-pNA; Sigma-Aldrich, St. Louis, MO, USA) was prepared as a 50 mM stock solution in ultrapure water. Human fibrinogen (FBN; Sigma-Aldrich) was prepared as a 10 mg/mL stock solution in TBS. Corn trypsin inhibitor (CTI), bovine factor Xa and human plasma were obtained from Haematologic Technologies LLC (Essex Junction, VT, USA). Commercial prothrombin time (PT) and activated partial thromboplastin time (aPTT) reagents were supplied by SEKISUI MEDICAL Co., Ltd. (Tokyo, Japan). All stock solutions were stored at −20°C and subjected to no more than one freeze-thaw cycle.

### 2.3 Coagulation assay using PT/aPTT reagents

Plasma clotting times were evaluated using PT and aPTT reagents in each fish species, as described previously (Matsui et al, 2026). Briefly, coagulation was initiated by adding 25 μL of PT or aPTT reagent together with 10 mM CaCl_2_ and either 1 mg/mL human FBN or 100 μM HD-FPR-pNA (all final concentrations) to 10 μL of plasma obtained from Japanese flounder, common carp, bullhead shark, and gray bichir. The reaction mixture was diluted with TBS to a final volume of 100 μL. Human plasma was included as a positive control. To assess coagulation, absorbance was measured at 415 nm every 15 s for a total of 80 measurements. Coagulation time was defined as the time required to reach 50% of the maximum absorbance. Each experiment was performed using plasma samples obtained from three independent individuals.

### 2.4 Inhibition of the intrinsic like-like coagulation by CTI

To examine the effects of inhibitors on plasma coagulation, fish plasma samples exhibiting intrinsic pathway-like activity, including those from Japanese flounder, common carp, and gray bichir, were treated with different concentrations of inhibitors, and their clotting times were measured. Briefly, coagulation was initiated by adding CaCl_2_, human FBN, and 25 μL of either PT or aPTT reagent to 10 μL of sodium citrate-treated plasma from Japanese flounder or gray bichir. The final concentrations of CaCl_2_ and human FBN were 10 mM and 1 mg/mL, respectively. CTI was added at final concentrations of 1, 10, or 25 μg/mL (71.4, 714, 1786 nM), and the total reaction volume was adjusted to 100 μL with TBS. For common carp plasma, 10 μL of plasma was combined with the same final concentrations of CaCl_2_ and human FBN, together with CTI at final concentrations of 50, 100, or 250 μg/mL (3.58, 7.14, 17.9 µM). Clotting time was determined as described in Section 2.3. Each experiment was independently performed using plasma from three individuals, and samples without CTI treatment served as the control.

### 2.5 Factor Xa bypass assay for CTI-inhibited coagulation

To determine whether CTI-induced inhibition of intrinsic pathway-like coagulation in Japanese flounder plasma could be rescued by the addition of factor Xa, 10 µL of citrated Japanese flounder plasma was supplemented with CaCl, human FBN, and an aPTT reagent at the same final concentrations as those described in Section 2.4. CTI was added at a final concentration of 25 µg/mL (1.79 µM), followed by human factor Xa at final concentrations of 1, 10, or 100 nM. Clotting times were measured as described in Section 2.3. Samples without factor Xa or CTI were included as controls. The experiment was performed using plasma samples independently collected from three individual fish.

### 2.6 Sequence acquisition and preparation

Protein sequences annotated as hepatocyte growth factor activator (HGFA) or factor XII in vertebrate species were retrieved from the NCBI Protein database (https://www.ncbi.nlm.nih.gov/protein/). The domain architecture of each protein sequence was predicted using ExPASy PROSITE (https://prosite.expasy.org/).

### 2.7 Complex structure prediction of CTI with fish HGFA

Complex structures of CTI (PDB ID: 1BEA) with the protease domains of flounder HGFA (accession no. XP_019950976.2) and common carp HGFA (NCBI accession no. XP_042584712) were predicted using AlphaFold 3 (https://alphafoldserver.com/). As a reference for comparison, a complex model of human factor XII (NP_000496) and CTI was also generated. For each prediction, the amino acid sequence corresponding to the protease domain of factor XII or HGFA and the mature CTI sequence derived from chain A were entered as separate protein chains at a 1:1 stoichiometric ratio. Default settings were used for multiple-sequence alignment generation and structural template searches. Five structural models were generated for each complex and ranked according to the ranking_score provided by AlphaFold Server. The top-ranked model was used for structural analysis. Predicted intermolecular contacts between CTI and factor XII or HGFA were analyzed using UCSF ChimeraX version 1.12.

### 2.8 Assessment of charged residue distribution in EGF-1 domain

In human factor XII, a cluster of basic residues in the first epidermal growth factor-like domain (EGF-1), comprising Lys73, Lys74, Lys76, and Lys81, forms an anion-binding site required for productive interactions with polyanionic substances such as polyphosphate (Shamanaev et al. 2023). To assess whether a corresponding basic patch is conserved in fish hepatocyte growth factor activator (HGFA), three-dimensional structural models were generated and analyzed. The amino acid sequences of HGFA from Japanese flounder, common carp, and California horn shark (*Heterodontus francisci*) were submitted to the AlphaFold 3 server to generate three-dimensional structural models. Basic amino acid residues (Lys, Arg, and His) and acidic amino acid residues (Asp and Glu) within the EGF-1 domain were displayed in different colors. The spatial clustering of basic residues and the potential formation of a basic surface corresponding to the Lys patch in human factor XII were visually assessed in each fish HGFA model.

### 2.9 Statistical analysis

Background fluorescence and absorbance were corrected by subtracting the signals obtained from the blank wells. In the inhibitor experiments using CTI, the inhibitor-treated group was compared with the corresponding inhibitor-free control group using one-way ANOVA, followed by Dunnett’s multiple-comparisons test. Statistical significance was defined as *p* < 0.05, with *p* < 0.05 and *p* < 0.001 indicated by one and three asterisks, respectively. For the bypass assay using human factor Xa, normality was assessed using the Shapiro– Wilk test. Because the assumption of normality was not satisfied, differences among the groups were analyzed using the Friedman test, followed by Conover’s post hoc test with Holm correction for multiple comparisons. Groups with statistically significant differences are indicated by different letters. Statistical analyses were performed using JASP version 0.18.3 (JASP Team).

## 3. Results

### 3.1 Clotting assay based on fibrin formation

Plasma coagulation was initiated using PT or aPTT reagents, and fibrin formation was evaluated by measuring turbidity. Following the addition of aPTT reagent, an increase in absorbance was observed in the plasma of teleost species, including common carp and Japanese flounder, as well as in human plasma used as a positive control (Figures 1B, D, and J). In contrast, no increase in absorbance was detected in gray bichir or bullhead shark plasma (Figures 1F and H). Following the addition of PT reagent, an increase in absorbance was observed in Japanese flounder, bullhead shark, and human plasma (Figure 1C, E, and I). However, the absorbance profile of bullhead shark plasma was highly variable, and the well-defined plateau observed in other species was not detected (Figure 1E). In gray bichir plasma, no PT reagent-induced increase in absorbance was detected within the observation period used in this study (Figures 1A and G).

**Figure 1.**
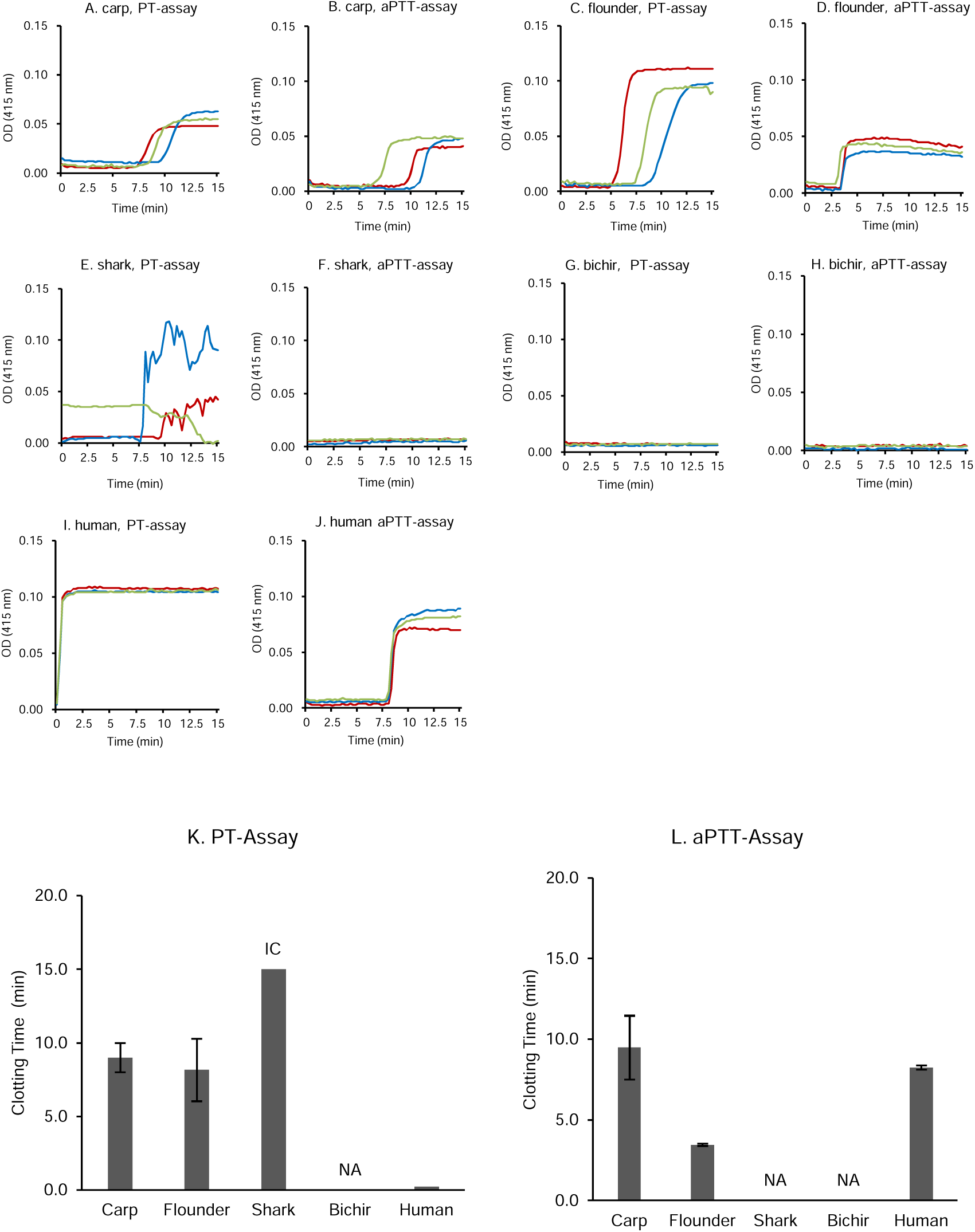
Clotting assay using fibrinogen. Prothrombin time (PT)-like and activated partial thromboplastin time (aPTT)-like plasma clotting activities were measured in several fish species using human fibrinogen as an exogenous substrate. Plasma from common carp (A, B), Japanese flounder (C, D), bullhead shark (E, F), and gray bichir (G, H) was mixed with PT or aPTT reagent at 25% (w/w), and coagulation was initiated by adding 10 mM CaCl and 1 mg/mL human fibrinogen at the indicated final concentrations. Absorbance at 415 nm was recorded every 15 s for up to 15 min. Clotting time was defined as the time required to reach half of the maximum absorbance (K, L). Human plasma served as a positive control (I, J). Results were designated as incomplete clotting (IC) when the measurement ended before the maximum absorbance was reached in at least one sample, and as no absorbance increase (NA) when at least one sample showed no marked increase in absorbance. Experiments were performed using plasma from three individual fish.

### 3. 2 Clotting assay based on thrombin substrate

The responsiveness of plasma from each fish species to PT and aPTT reagents was also evaluated based on thrombin-like enzymatic activity. Following the addition of aPTT reagent, the absorbance of Japanese flounder and gray bichir plasma increased and reached a clear plateau (Figures 2D and H). Human plasma also showed an increase in absorbance, although the response did not reach a plateau within the observation period. In contrast, no increase in absorbance was detected in common carp or bullhead shark plasma (Figures 2B and F). Following the addition of PT reagent, an increase in absorbance was observed in plasma from Japanese flounder, bullhead shark, gray bichir, and human (Figures 2C, E, G, and I), whereas no increase was detected in common carp plasma (Figure 2A). As observed in the fibrin formation-based assay, the absorbance profile of bullhead shark plasma was noisy (Figure 2E).

**Figure 2.**
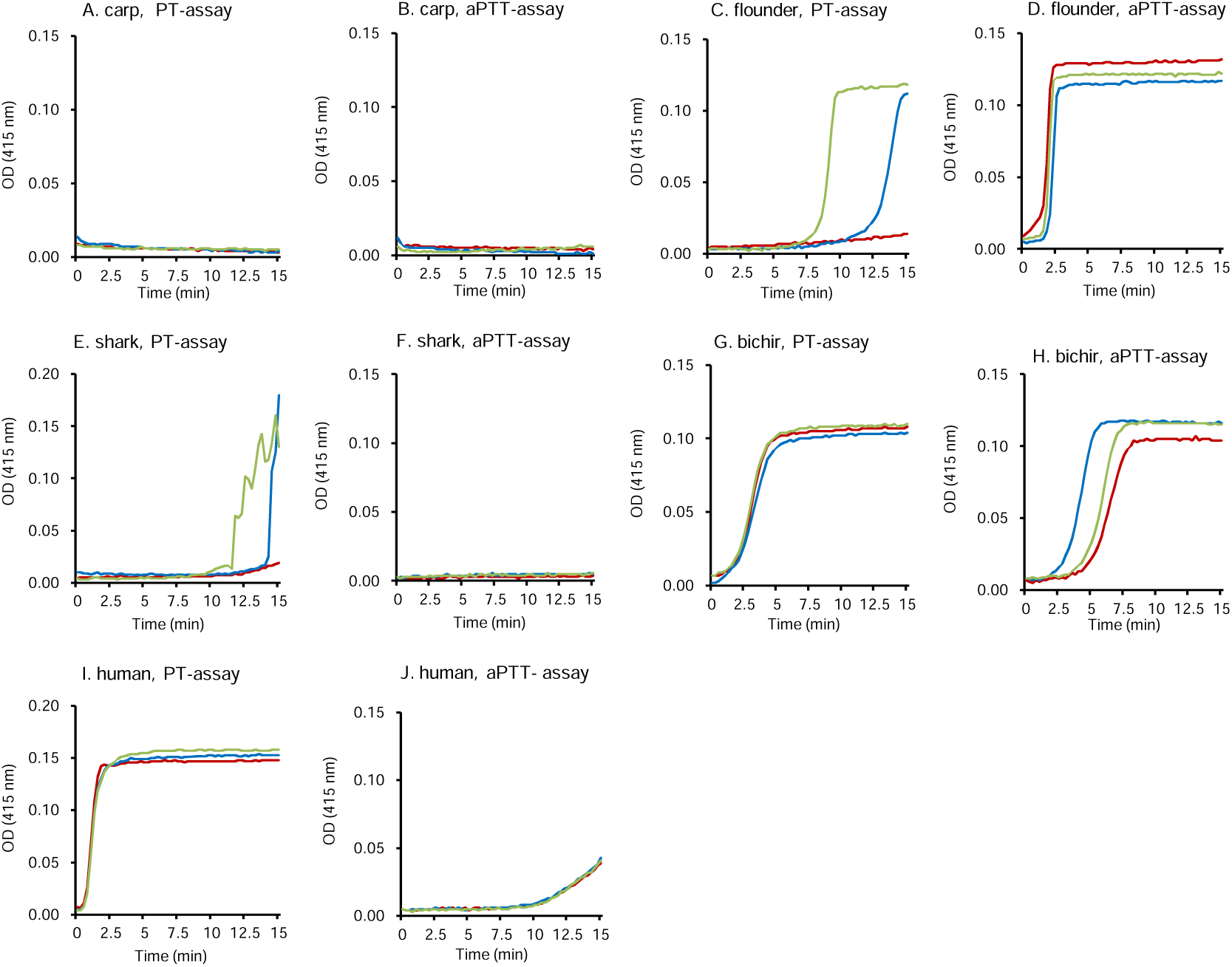
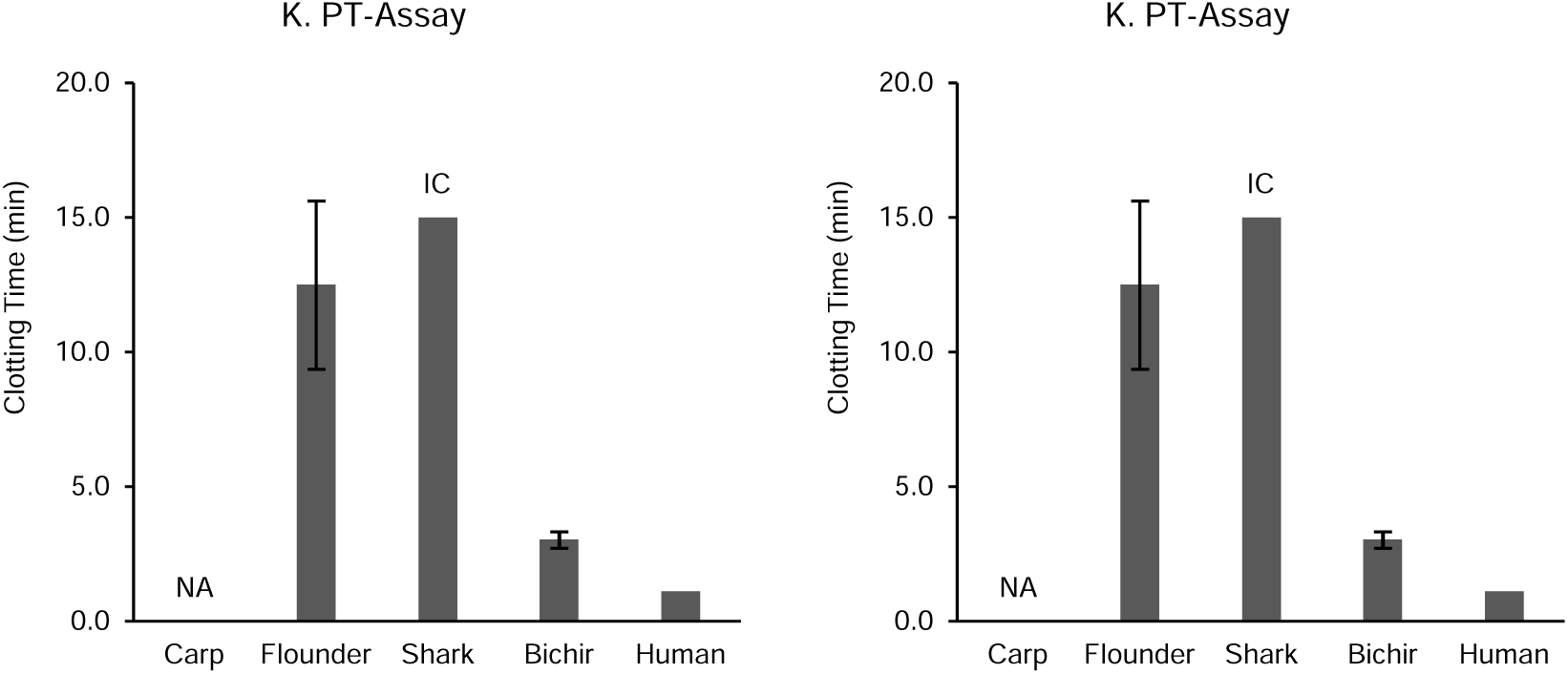
Thrombin substrate assay. Plasma clotting times in several fish species were measured using prothrombin time (PT) and activated partial thromboplastin time (aPTT) reagents with the thrombin substrate Nα-(2-hydroxy-3,4-difluorophenylacetyl)-D-phenylalanyl-L-arginine *p*-nitroanilide (HD-FPR-pNA). Plasma from common carp (A, B), Japanese flounder (C, D), bullhead shark (E, F), and gray bichir (G, H) was mixed with PT or aPTT reagent at 25% (w/w). Coagulation was initiated by adding CaCl and HD-FPR-pNA at final concentrations of 10 mM and 100 μM, respectively. Absorbance was measured at 415 nm every 15 s for up to 15 min. Clotting time was defined as the time required for the absorbance to reach half of its maximum value (K, L). Human plasma was used as a positive control (I, J). When the measurement ended before the absorbance reached its maximum in at least one sample, the result was designated as incomplete clotting (IC). When no marked increase in absorbance was observed in at least one sample, the result was designated as no absorbance increase (NA). Experiments were performed using plasma samples from three independent fish per species.

### 3.3 Effects of CTI on intrinsic-like blood coagulation

Plasma from common carp, Japanese flounder, and gray bichir was activated with aPTT reagent to evaluate the inhibitory effect of CTI. In Japanese flounder plasma, treatment with CTI at 10 and 25 μg/mL significantly prolonged the clotting time compared with that in the absence of CTI (Figure 3B). In contrast, CTI treatment did not significantly alter the clotting time at any concentration tested under PT-like assay conditions using Japanese flounder plasma (Figure 3A). Under aPTT-like assay conditions, the coagulation time of common carp plasma was significantly prolonged relative to that in the absence of CTI only following treatment with 250 μg/mL CTI, a concentration tenfold higher than the maximum effective concentration used for Japanese flounder plasma (Figure 3D). CTI-induced prolongation of the clotting time was also observed in common carp plasma under PT-like assay conditions (Figure 3C). In gray bichir plasma, CTI treatment at concentrations up to 25 μg/mL did not significantly prolong the coagulation time under either PT- or aPTT-like assay conditions (Figures 3E and F).

**Figure 3.**
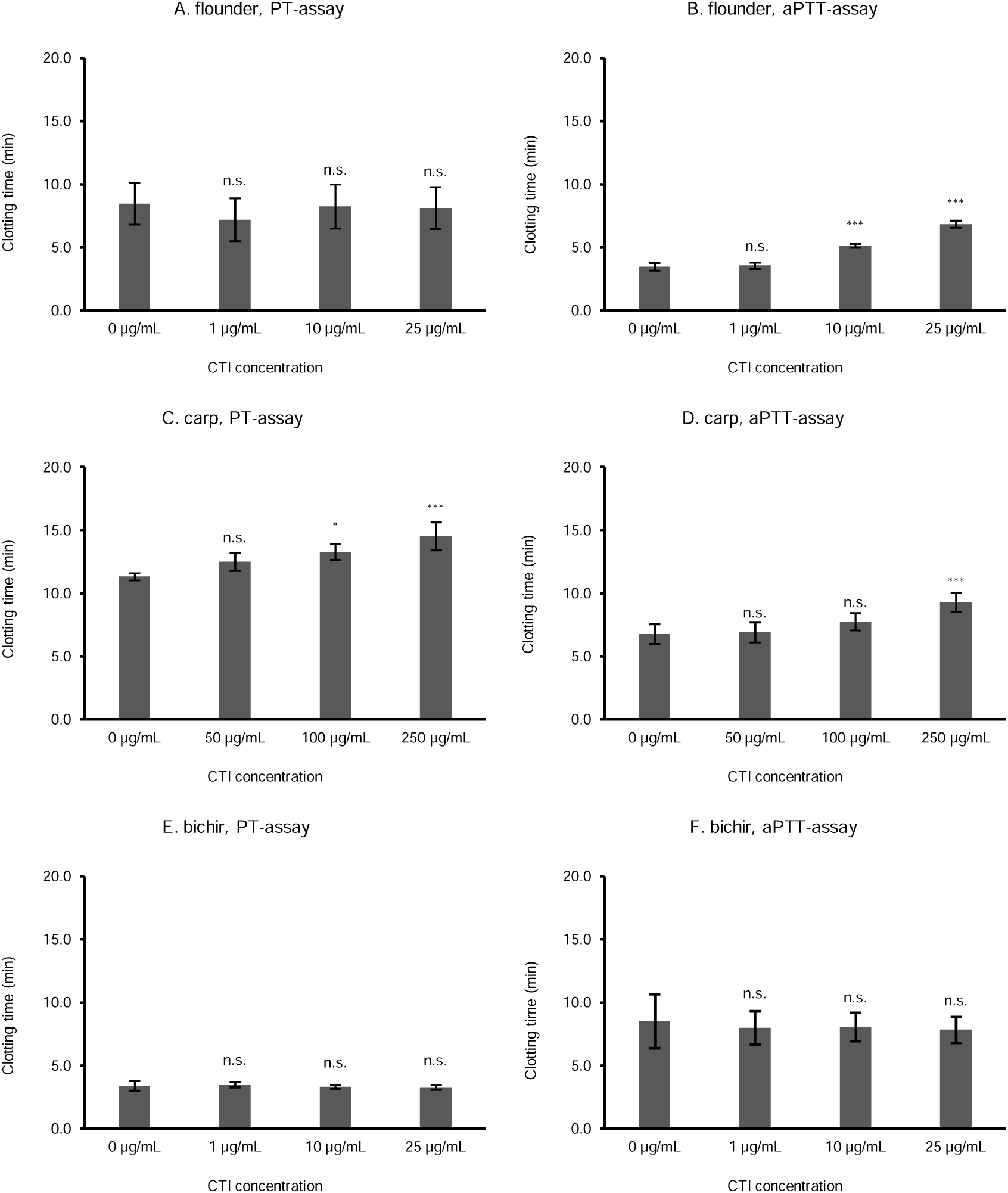
factor XII inhibitor on fish contact coagulation. The effects of corn trypsin inhibitor (CTI), an inhibitor of human factor XIIa, on intrinsic pathway-like coagulation activity were investigated in plasma from Japanese flounder, common carp, and gray bichir. Ten microliters of citrated plasma from each fish species was mixed with 10 μL of a solution containing CaCl and human fibrinogen at final concentrations of 10 mM and 1 mg/mL, respectively. CTI was added at final concentrations of 1, 10, and 25 μg/mL (71.4, 714, 1786 nM) for Japanese flounder and gray bichir plasma and 50, 100, and 250 μg/mL (3.58, 7.14, 17.9 µM) for common carp plasma. Plasma coagulation was activated by adding 25 μL of PT (A, C, E) or aPTT (B, D, F) reagents. Absorbance at 415 nm was measured every 15 s, and clotting time was defined as the time required for the absorbance to reach 50% of its maximum value. Clotting times at each CTI concentration were compared with those in the absence of CTI using one-way ANOVA, followed by Dunnett’s multiple-comparison test. Experiments were performed using three independent plasma samples from each fish species (n = 3). *** *p* < 0.01, * *p* < 0.05, n.s., *p* > 0.05.

### 3.4 Factor Xa bypass assay for CTI-inhibited coagulation

Japanese flounder plasma activated with an aPTT reagent was treated with CTI, and human factor Xa was subsequently added to determine whether the prolonged clotting time could be restored to the level observed in the absence of CTI. In CTI-treated plasma, the addition of human factor Xa significantly shortened clotting time (Figure 4A). The clotting times did not differ significantly between the groups without factor Xa and with 1 nM factor Xa. In contrast, the addition of 10 or 100 nM factor Xa significantly shortened the clotting time compared to the group without factor Xa (Figure 4B).

**Figure 4.**
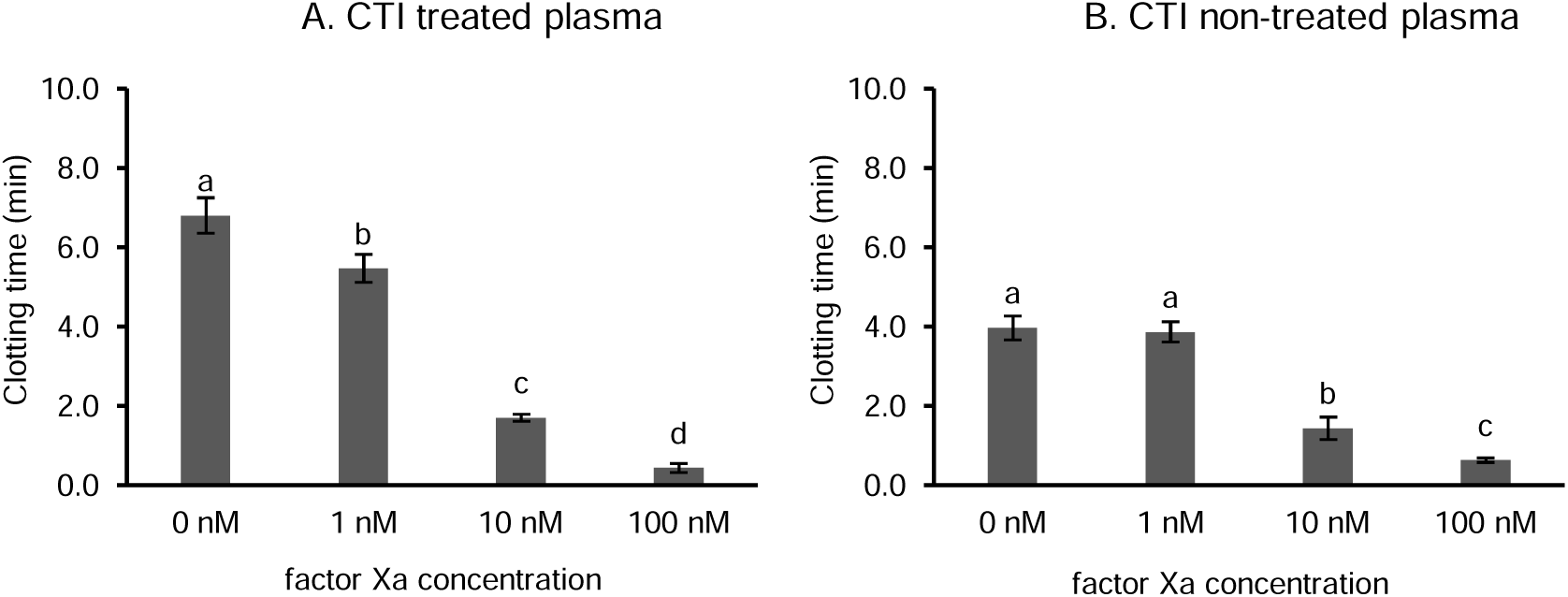
factor Xa bypass from CTI Inhibition. Restoration of corn trypsin inhibitor (CTI)-inhibited activated partial thromboplastin time (aPTT)-like coagulation activity in Japanese flounder plasma by human factor Xa. We examined whether the addition of human factor Xa restored aPTT-like coagulation activity in Japanese flounder plasma following inhibition of contact activation by CTI. A reaction mixture containing 10 μL of 100 mM CaCl, 10 μL of 25 μg/mL CTI, 25 μL of activated partial thromboplastin time (aPTT) reagent, 10 μL of 10 mg/mL human fibrinogen, and 10 μL of human factor Xa at 10, 100, or 1000 nM was added to 10 μL of citrated Japanese flounder plasma to initiate coagulation. Absorbance was measured at 415 nm every 15 s, and clotting time was defined as the time required for the absorbance to reach 50% of its maximum value. Reactions were performed in the presence of CTI (A) or an equivalent volume of Tris buffer instead of CTI (B). Experiments were performed using three independent plasma samples (n = 3). Normality was assessed using the Shapiro–Wilk test. Differences among factor Xa concentrations were subsequently analyzed using the Friedman test, followed by Conover’s post hoc test with Holm correction for multiple comparisons. Different letters indicate significant differences (*p* < 0.05).

### 3.5 Prediction of flounder and carp HGFA–CTI complexes

To compare the structural features potentially associated with the difference in CTI sensitivity between the intrinsic pathway-like activities of carp and Japanese flounder, interactions in these fish HGFA–CTI complex models were analyzed with reference to the previously reported human factor XIIa–CTI complex model (Hamad et al., 2017). In the human factor XIIa–CTI complex model, a salt bridge was predicted between the NH2 atom of the CTI Arg34 side chain and the OD2 atom of the factor XIIa Asp189 side chain (Figure 5C). In the fish HGFA–CTI complex models, the NH2 atom of CTI Arg34 was also positioned close to the OD2 atom of the residue corresponding to factor XIIa Asp189, with interatomic distances of 2.773 and 2.740 Å, respectively (Figures 5A, B). In addition, the aromatic rings of CTI Trp22 and factor XIIa Tyr99 formed a parallel-displaced π–π stacking interaction in the human factor XIIa–CTI complex model (Figure 5F). The residues corresponding to factor XIIa Tyr99 in carp and Japanese flounder HGFA did not possess aromatic side chains, and no corresponding π–π stacking interactions were observed in either HGFA–CTI complex model (Figure 5D, E). A hydrogen bond between the NE1 atom of CTI Trp22 and the backbone of factor XIIa Pro96 was also predicted in the human factor XIIa–CTI complex model (Figure 5F). Although Pro86 and Pro87 occupied the corresponding positions in carp and Japanese flounder HGFA, respectively, no corresponding hydrogen bond was identified in either HGFA–CTI complex model (Figure 5D, E). Furthermore, the human factor XIIa–CTI complex model contains a salt bridge between CTI Arg43 and factor XIIa Asp60A (Figure 5I). In the Japanese flounder HGFA–CTI complex model, the distance between the NE atom of CTI Arg43 and the OD2 atom of Asp24, corresponding to factor XIIa Asp60A, was 4.120 Å (Figure G). In contrast, in the carp HGFA–CTI complex model, the CTI Arg43 side chain was not positioned close to the side chain of Asp22, which corresponds to factor XIIa Asp60A (Figure 5I).

**Figure 5.**
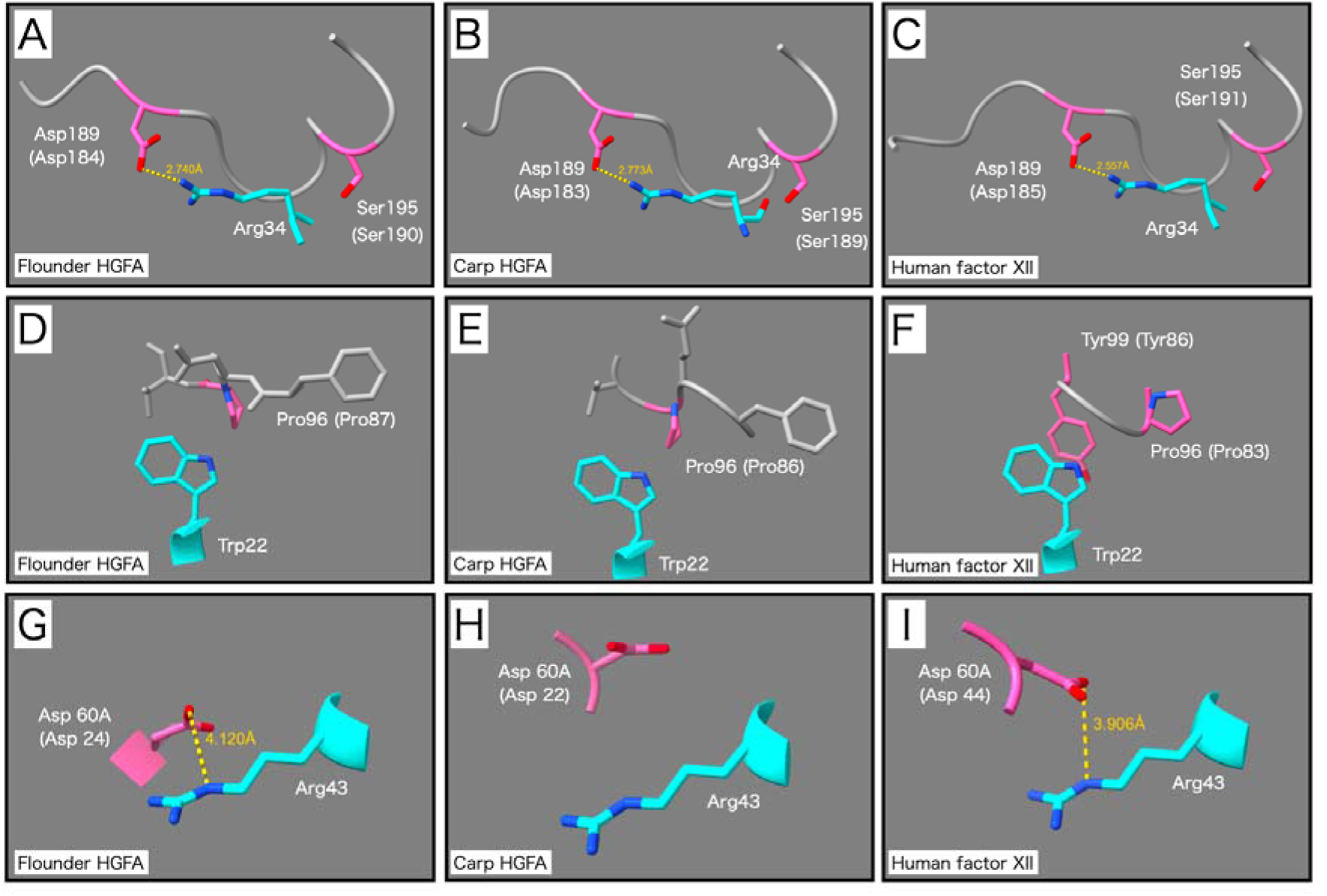
Predicted binding of CTI to human factor XII and fish HGFAs. Predicted interactions of corn trypsin inhibitor (CTI) with human factor XII and fish hepatocyte growth factor activators (HGFAs). Complex models of CTI (PDB ID: 1BEA) with trypsin-like serine protease domains of human factor XII (accession no. NP_000496), Japanese flounder HGFA (accession no. XP_019950976.2), and common carp HGFA (accession no. XP_042584712) were generated using AlphaFold 3 to assess their structural compatibility with CTI binding. Potential salt-bridge formation between CTI Arg34 and protease Asp189 was examined in the human factor XII–CTI, Japanese flounder HGFA–CTI, and common carp HGFA–CTI models (A–C, respectively). Potential interactions between CTI Trp22 and protease Pro96 (D–F) and between CTI Arg43 and protease Asp60A (G–I) were examined in the same order. Catalytically important protease residues are shown in hot pink, and the inhibitory region of CTI is shown in cyan. The yellow and light-blue dashed lines represent the potential salt bridges and predicted hydrogen bonds, respectively. Protease residues were numbered according to the chymotrypsin numbering system, with the corresponding residue numbers in each protein shown in parentheses.

### 3.6 Assessment of charged residue distribution in the first EGF domain

The potential structural conservation of the cluster of basic residues in the EGF-1 domain involved in the polyanion-dependent autoactivation of human factor XII was examined in fish HGFA. In the AlphaFold-predicted structure of human factor XII, the EGF-1 Lys cluster comprising Lys73, Lys74, Lys76, and Lys81 was reproduced on the same molecular surface (Figure 6D). In contrast, no distinct clustering of basic residues was observed in the corresponding N-terminal region of the EGF-1 domain of fish HGFA (Figure 6A-C). Similarly, no basic surface corresponding to the Lys cluster of human factor XII was predicted in the N-terminal region of the EGF-1 domain of Japanese flounder, common carp, or California horn shark HGFA (Figures 6A–C). In each fish HGFA, a region containing two or more closely positioned Lys and/or Arg residues was observed on the C-terminal side of the EGF-1 domain; however, acidic residues were also present in the vicinity of these basic

**Figure 6.**
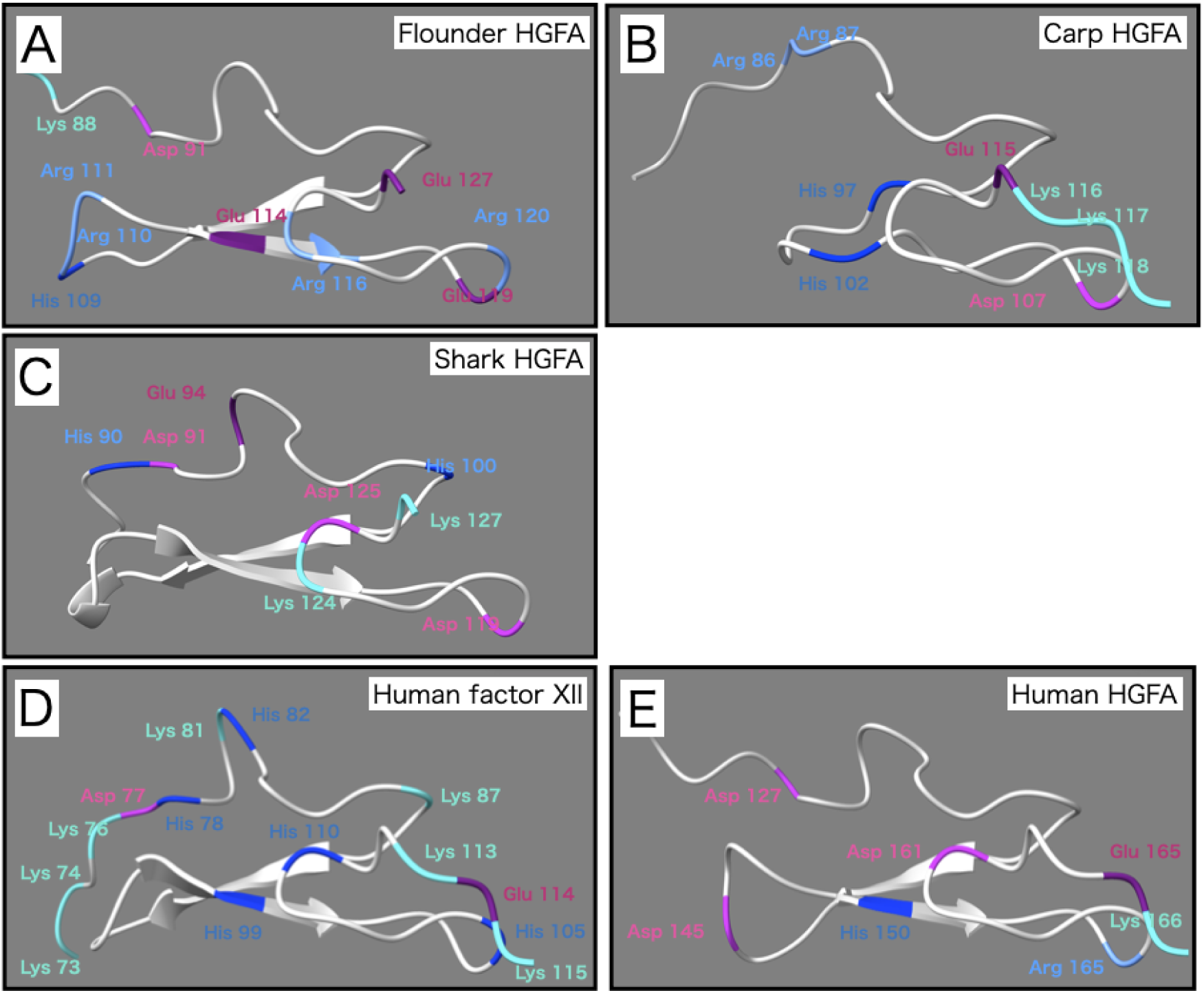
Structural analysis of EGF-1 domains using three-dimensional modeling. The first epidermal growth factor-like domain (EGF-1 domain) of human factor XII contains a surface-exposed Lys cluster that mediates interactions with negatively charged surfaces and promotes surface-dependent autoactivation. To determine whether the EGF-1 domains of fish hepatocyte growth factor activators (HGFAs) contain similar clusters of basic residues, three-dimensional models of Japanese flounder HGFA (A; accession no. XP_019950976.2), common carp HGFA (B; accession no. XP_042584712), and California horn shark HGFA (C; accession no. XP_067885357.1) were generated using AlphaFold 3, and the EGF-1 domains were extracted for analysis. Human factor XII (D; accession no. NP_000496) and human HGFA (E; accession no. Q04756) were analyzed in the same manner as controls. The structures were visualized using UCSF ChimeraX. Lys, Arg, His, Asp, and Glu residues are shown in cyan, forest green, blue, magenta, and purple, respectively.

## Discussion

Intrinsic pathway-like coagulation activity was detected in multiple ray-finned fish lineages; however, its detection depended partly on the assay substrate, and its sensitivity to CTI varied substantially among species. Furthermore, the occurrence of aPTT-like activity did not consistently correspond to the genomic presence or domain architecture of HGFA, its predicted ability to bind CTI, or the presence of structural features associated with factor XII-like contact activation. These findings suggest that the mechanisms initiating intrinsic pathway-like coagulation have diverged among fish species, and that the HGFA-mediated mechanism proposed in zebrafish should not be regarded as universally conserved in fishes.

In the teleosts common carp and Japanese flounder, coagulation responses to the aPTT reagent were detected using a human fibrinogen-based assay (Figure 1). Although previous studies have used different reagents and assay conditions, aPTT-based clotting has also been measured in zebrafish, Japanese flounder, and common carp (Kawatsu 1986; Dhinoja et al. 2024; Matsui 2026), consistent with the presence of intrinsic pathway-like coagulation capacity in these species. In the thrombin-substrate-based assay, however, both PT and aPTT reagents induced substrate-cleaving activity in Japanese flounder plasma, whereas neither reagent produced a detectable response in common carp plasma (Figures 2A–D). Given that coagulation was detected in the fibrinogen-based assay, the lack of an increase in absorbance with the thrombin substrate may reflect differences in substrate recognition between fish thrombins rather than the absence of thrombin generation. Instead, structural differences between fish thrombins may result in different substrate recognitions.

In bichir plasma, thrombin-substrate-cleaving activity was detected under aPTT-like conditions, suggesting the presence of an intrinsic pathway-like mechanism capable of generating thrombin-like protease activity in this early-diverging ray-finned fish lineage (Figures 2G and 2H). In contrast, no increase in turbidity was detected in the fibrinogen-based assay (Figures 1G and 1H). Because the same assay detected fibrin formation in common carp, Japanese flounder, shark, and human plasma, the negative result in bichir is unlikely to reflect a general failure of the assay. Instead, thrombin-like protease activity may have been generated but failed to efficiently cleave human fibrinogen. Alternatively, human fibrinogen may have been cleaved, but the resulting fibrin monomers may not have undergone efficient protofibril assembly, lateral aggregation, or fiber formation, all of which contribute to clot turbidity (Pieters and Wolberg 2019). In either case, the absence of turbidity does not necessarily indicate a lack of thrombin-like protease activity, but rather suggests limited functional compatibility between the bichir coagulation system and human fibrinogen.

In shark plasma, no aPTT-like response was detected within the measurement period using either detection system, whereas PT-like reactions were observed, although the resulting absorbance curves were unstable (Figures 1E and 1F; Figure 2E and 2F). Rapid dissolution of newly formed fibrin clots has been previously reported in sharks (Doolittle and Surgenor 1962; Doolittle 1963). Thus, the irregular absorbance changes observed here may reflect fibrin formation in response to the PT reagent, followed by rapid clot dissolution and a heterogeneous redistribution.

Taken together, the observed functional differences did not correspond directly to the presence of HGFA. aPTT-induced thrombin-substrate-cleaving activity was detected not only in Japanese flounder and common carp, which possess canonical HGFA orthologs, but also in bichir, in which no clear HGFA ortholog has been identified (Figures 1 and 2; Chana-Muñoz et al. 2019). Conversely, sharks possess an HGFA-like molecule with an atypical domain duplication but showed no detectable aPTT-like activity under the present conditions (Figures 1). Therefore, HGFA sequence information alone cannot predict the occurrence of aPTT-like coagulation activity in fish.

Commercial clinical coagulation reagents have been used to determine plasma clotting times in several fish species (Fujikata and Ikeda, 1985; Kawatsu, 1986; Iyer and Jagadeeswaran, 2021), but the resulting values should not be treated as universal measures of coagulation capacity. Fish coagulation factors may differ in their interactions with reagent components, including tissue factor, phospholipids, and contact activators, owing to interspecific differences in their primary structures and molecular interactions (Doolittle and Surgenor 1962; Sheehan et al. 2001). Moreover, even in clinical laboratories, clotting times vary according to the reagents, instruments, and reaction conditions used (Gardiner et al. 2021; Kato et al. 2022). Therefore, direct comparison with previously reported fish clotting times obtained under different conditions is difficult. Nevertheless, because the species examined here were evaluated under identical conditions, the present assays provide useful relative indices for their responses to the same coagulation stimuli and substrates.

CTI inhibition experiments further revealed substantial interspecific differences in the intrinsic pathway-like coagulation. In Japanese flounder, CTI prolonged aPTT-like clotting at concentrations comparable to those used to suppress contact activation in mammalian blood, while having no significant effect on PT-like clotting (Figures 3A and 3B; Butenas and Mann, 2004). This pattern is consistent with the selective inhibition of an intrinsic pathway-like reaction rather than the nonspecific inhibition of the common pathway. Furthermore, exogenous human factor Xa restored clotting in the presence of CTI, with 10 nM factor Xa reducing the clotting time to a level comparable to or shorter than that observed in the absence of CTI (Figure 4). As factor Xa functions downstream of the convergence of the intrinsic and extrinsic pathways and promotes thrombin generation as part of the prothrombinase complex (Schreuder et al. 2019), this restoration suggests that the primary CTI-sensitive step in Japanese flounder lies upstream of factor Xa generation, at least within the framework of the established coagulation cascade.

In contrast, aPTT-like activity in common carp was inhibited only at a high CTI concentration that also prolonged PT-like clotting (Figures 3C and 3D). Therefore, the effect observed in common carp cannot be readily interpreted as selective inhibition of the intrinsic pathway-like reaction. At high concentrations, CTI may inhibit a component shared by the PT-like and aPTT-like reactions or exert nonspecific effects on coagulation. Bichir plasma showed no detectable inhibition in either assay within the concentration range examined (Figures 3E and 3F), indicating lower CTI sensitivity than that observed in Japanese flounder and common carp.

Structural modeling did not fully explain the interspecific difference in CTI sensitivity. In both Japanese flounder and common carp HGFA models, the reactive-site loop of CTI could be accommodated within the catalytic site, and the positions of CTI Arg34 and HGFA Asp189 were compatible with a canonical interaction (Figures 5A–C). However, the interactions involving the S3 pocket that contribute to the specific binding of CTI to human factor XIIa were not reproduced in either fish HGFA model. In particular, Tyr99 of human factor XIIa (Tyr86 according to the mature-protein numbering), which contributes to the interaction with CTI, was not conserved in either fish HGFA. Moreover, no alternative aromatic side chain was positioned to reproduce the corresponding interaction with CTI (Figures 5D–F; Hamad et al. 2017). The predicted salt-bridge interaction between Asp60A of HGFA and Arg43 of CTI differed between Japanese flounder and common carp because of the distinct orientation of the Asp60A side chain (Figures 5G–I). Although these differences between human factor XIIa and fish HGFAs may affect CTI-binding affinity to some extent, they do not preclude the fundamental capacity of fish HGFAs to bind CTI. Thus, the predicted HGFA–CTI interactions alone cannot account for the marked difference in plasma CTI sensitivity between Japanese flounder and common carp. Other CTI-sensitive proteases, species-specific regulatory factors, or differences in the relative contribution of HGFA to plasma aPTT-like activity may also be involved.

The modeled EGF-1 domains also lacked the major structural feature required for factor XII-like contact activation. In human factor XII, a cluster comprising Lys73, Lys74, Lys76, and Lys81 mediates productive interactions with polyanionic surfaces and subsequent surface-dependent activation (Shamanaev et al. 2022, 2023). The corresponding lysine cluster is absent from human HGFA, and the replacement of the factor XII EGF-1 domain with the corresponding HGFA sequence reportedly abolishes procoagulant activity (Shamanaev et al. 2022). Consistent with these observations, none of the fish HGFA models showed a comparable cluster of basic residues on the N-terminal side of the EGF-1 domain (Figures 6A–C). Although basic residues were present elsewhere in the domain, they were dispersed and accompanied by acidic residues, making the formation of a strongly positively charged surface improbable. These structural features suggest that fish HGFA is unlikely to undergo EGF-1-dependent autoactivation on negatively charged surfaces via the same mechanism as human factor XII.

Although flounder plasma showed CTI-sensitive coagulation activity (Figures 3–5), structural analysis of flounder HGFA did not support factor XII-like, surface-dependent autoactivation (Figure 6); however, these findings are not necessarily contradictory. A previous study in zebrafish demonstrated changes in coagulation assays following HGFA knockdown or inhibitor treatment but did not establish that HGFA directly initiates coagulation (Alharbi et al., 2021). If HGFA does not function as an initiator of coagulation but rather in the amplification or propagation of coagulation initiated through another mechanism, the CTI sensitivity of olive flounder plasma can be reconciled with the structural findings for HGFA.

In conclusion, intrinsic pathway-like coagulation activity is present in several fish lineages, including an early-diverging ray-finned fish, but its detection and inhibitor sensitivity differ among species. The occurrence of this activity did not consistently correlate with the HGFA sequence architecture, predicted CTI binding, or structural features associated with factor XII-like contact activation. HGFA may contribute to intrinsic-like coagulation in particular fish species, including Japanese flounder, but the present findings do not support its uniform designation as a common factor of intrinsic pathway-like coagulation across fishes. Instead, the initiation mechanisms and relative contributions of CTI-sensitive proteases appear to have diversified during fish evolution.

## CRediT authorship contribution statement

**Shintaro Matsui:** Conceptualization, Methodology, Validation, Investigation, Formal analysis, Resources, Data curation, Visualization, Project administration, Software, Writing Original Draft, Funding acquisition. **Asami Yoshida:** Data curation, Writing – review & editing.

## Data availability

Data are available upon reasonable request.

## Acknowledgements

The authors thank Dr. Takafumi Oyoshi for providing valuable advice on the construction of the protein complex models.

## Declaration of generative AI and AI-assisted technologies in the manuscript preparation process

During the preparation of this manuscript, the author used Copilot for English editing. After using this tool, the author reviewed and edited the content as needed and took full responsibility for the content of the published article.

## References

Alharbi, A., Iyer, N., Al Qaryoute, A., Raman, R., Burks, D.J., Azad, R.K., Jagadeeswaran, P., 2021. Role of ribosomal RNA released from red cells in blood coagulation in zebrafish and humans. Blood Adv. 5, 4634– 4647. 10.1182/bloodadvances.2020003325.

Beeler, D.L., Aird, W.C., Grant, M.A., 2018. Evolutionary conservation of the allosteric activation of factor VIIa by tissue factor in lamprey. J. Thromb. Haemost. 16, 734–748. 10.1111/jth.13968.

Butenas, S., Mann, K.G., 2004. Active tissue factor in blood? Nat. Med. 10, 1155–1156. 10.1038/nm1104-1155b.

Chana-Muñoz, A., Jendroszek, A., Sønnichsen, M., Wang, T., Ploug, M., Jensen, J.K., Andreasen, P.A., Bendixen, C., Panitz, F., 2019. Origin and diversification of the plasminogen activation system among chordates. BMC Evol. Biol. 19, 27. 10.1186/s12862-019-1353-z.

Davie, E.W., Ratnoff, O.D., 1964. Waterfall sequence for intrinsic blood clotting. Science 145, 1310–1312. 10.1126/science.145.3638.1310.

Dhinoja, S., De Maria, A., Al Qaryoute, A., Jagadeeswaran, P., 2024. Characterization of zebrafish coagulation cofactors Fviii and Fv mutants and modeling hemophilia A and factor V deficiency. Blood Coagul. Fibrinolysis 35, 238–247. 10.1097/MBC.0000000000001308.

Fujikata, A., Ikeda, Y., 1985. Blood coagulation and clotting tests in carp. Nippon Suisan Gakkaishi 51, 933– 939.

Fukushima, T., Uchiyama, S., Tanaka, H., Kataoka, H., 2018. Hepatocyte growth factor activator: a proteinase linking tissue injury with repair. Int. J. Mol. Sci. 19, 3435. 10.3390/ijms19113435.

Gardiner, C., Coleman, R., de Maat, M.P.M., Dorgalaleh, A., Echenagucia, M., Gosselin, R.C., Ieko, M., Kitchen, S., 2021. International Council for Standardization in Haematology laboratory guidance for the evaluation of haemostasis analyser–reagent test systems. Part 1: Instrument-specific issues and commonly used coagulation screening tests. Int. J. Lab. Hematol. 43, 169–183. 10.1111/ijlh.13411.

Hanumanthaiah, R., Day, K., Jagadeeswaran, P., 2002. Comprehensive analysis of blood coagulation pathways in Teleostei: evolution of coagulation factor genes and identification of zebrafish factor VIIi. Blood Cells Mol. Dis. 29, 57–68. 10.1006/bcmd.2002.0534.

Hamad, B.K., Pathak, M., Manna, R., Fischer, P.M., Emsley, J., Dekker, L.V., 2017. Assessment of the protein interaction between coagulation factor XII and corn trypsin inhibitor by molecular docking and biochemical validation. J. Thromb. Haemost. 15, 1818–1828. 10.1111/jth.13773.

Hoffman, M., Monroe, D.M., 2001. A cell-based model of hemostasis. Thromb. Haemost. 85, 958–965. 10.1055/S-0037-1615947.

Iyer, N., Jagadeeswaran, P., 2021. Microkinetic coagulation assays for human and zebrafish plasma. Blood Coagul. Fibrinolysis 32, 50–56. 10.1097/MBC.0000000000000975.

Jiang, Y., Doolittle, R.F., 2003. The evolution of vertebrate blood coagulation as viewed from a comparison of puffer fish and sea squirt genomes. Proc. Natl. Acad. Sci. U.S.A. 100, 7527–7532. 10.1073/pnas.0932632100.

Kannemeier, C., Shibamiya, A., Nakazawa, F., et al., 2007. Extracellular RNA constitutes a natural procoagulant cofactor in blood coagulation. Proc. Natl. Acad. Sci. U.S.A. 104, 6388–6393. 10.1073/pnas.0608647104.

Kataoka, H., Kawaguchi, M., 2010. Hepatocyte growth factor activator (HGFA): pathophysiological functions in vivo. FEBS J. 277, 2230–2237. 10.1111/j.1742-4658.2010.07640.x.

Kato, K., Hatayama, Y., Shimohiro, H., Ichikawa, H., Fukuda, T., 2022. Differences in the composition of activated partial thromboplastin time reagents affect clot waveform analysis. Yonago Acta Med. 65, 226–234. 10.33160/yam.2022.08.013.

Kawatsu, H., 1986. Clotting time of common carp blood. Nippon Suisan Gakkaishi 52, 591–595. 10.2331/suisan.52.591.

Long, A.T., Kenne, E., Jung, R., Fuchs, T.A., Renné, T., 2016. Contact system revisited: an interface between inflammation, coagulation, and innate immunity. J. Thromb. Haemost. 14, 427–437. 10.1111/jth.13235.

Macfarlane, R.G., 1964. An enzyme cascade in the blood clotting mechanism, and its function as a biochemical amplifier. Nature 202, 498–499. 10.1038/202498a0.

Matsui, S., 2026. Dual effects of mucus from the Japanese flounder (*Paralichthys olivaceus*) on blood coagulation: procoagulant and anticoagulant activities. Comp. Biochem. Physiol. B Biochem. Mol. Biol. 111285. 10.1016/j.cbpb.2026.111285.

Matsui, S., Nakamura, O., Nakao, M., Oyoshi, T., Tsutsui, S., 2026. A non-enzymatic C-type lectin from bullhead shark Heterodontus japonicus skin amplifies coagulation via the common pathway downstream of factor Xa. Biochimie, in press. 10.1016/j.biochi.2026.10.023.

Matsui, S., Nakamura, O., Tsutsui, S., 2023. Unique properties of prothrombin in the bullhead shark *Heterodontus japonicus*: the first report of purification and characterization of a blood coagulation factor in Chondrichthyes. J. Comp. Physiol. B 193, 71–80. 10.1007/s00360-022-01472-3.

Miyazawa, K., Shimomura, T., Kitamura, N., 1996. Activation of hepatocyte growth factor in the injured tissues is mediated by hepatocyte growth factor activator. J. Biol. Chem. 271, 3615–3618. 10.1074/jbc.271.7.3615.

Mukai, S., Fukushima, T., Naka, D., Tanaka, H., Osada, Y., Kataoka, H., 2008. Activation of hepatocyte growth factor activator zymogen (pro-HGFA) by human kallikrein 1-related peptidases. FEBS J. 275, 1003– 1017. 10.1111/j.1742-4658.2008.06265.x.

Müller, F., Mutch, N.J., Schenk, W.A., et al., 2009. Platelet polyphosphates are proinflammatory and procoagulant mediators in vivo. Cell 139, 1143–1156. 10.1016/j.cell.2009.11.001.

Park, S., Park, J.K., 2024. Back to basics: the coagulation pathway. Blood Res. 59, 35. 10.1007/s44313-024-00040-8.

Pathak, M., Kaira, B.G., Slater, A., Emsley, J., 2018. Cell receptor and cofactor interactions of the contact activation system and factor XI. Front. Med. 5, 66. 10.3389/fmed.2018.00066.

Pieters, M., Wolberg, A.S., 2019. Fibrinogen and fibrin: an illustrated review. Res. Pract. Thromb. Haemost. 3, 161–172. 10.1002/rth2.12191.

Ponczek, M.B., Gailani, D., Doolittle, R.F., 2008. Evolution of the contact phase of vertebrate blood coagulation. J. Thromb. Haemost. 6, 1876–1883. 10.1111/j.1538-7836.2008.03143.x.

Ponczek, M.B., Shamanaev, A., LaPlace, A., Dickeson, S.K., Srivastava, P., Sun, M.F., Gruber, A., Kastrup, C., Emsley, J., Gailani, D., 2020. The evolution of factor XI and the kallikrein-kinin system. Blood Adv. 4, 6135–6147. 10.1182/bloodadvances.2020002456.

Schreuder, M., Reitsma, P.H., Bos, M.H.A., 2019. Blood coagulation factor Va’s key interactive residues and regions for prothrombinase assembly and prothrombin binding. J. Thromb. Haemost. 17, 1229–1239. 10.1111/jth.14487.

Shamanaev, A., Ivanov, I., Sun, M.F., Litvak, M., Srivastava, P., Mohammed, B.M., Shaban, R., Maddur, A., Verhamme, I.M., McCarty, O.J.T., Law, R.H.P., Gailani, D., 2022. Model for surface-dependent factor XII activation: the roles of factor XII heavy chain domains. Blood Adv. 6, 3142–3154. 10.1182/bloodadvances.2021005976.

Shamanaev, A., Litvak, M., Cheng, Q., Ponczek, M., Dickeson, S.K., Smith, S.A., Morrissey, J.H., Gailani, D., 2023. A site on factor XII required for productive interactions with polyphosphate. J. Thromb. Haemost. 21, 1567–1579. 10.1016/j.jtha.2023.02.014.

Sheehan, J., Templer, M., Gregory, M., Hanumanthaiah, R., Troyer, D., Phan, T., Thankavel, B., Jagadeeswaran, P., 2001. Demonstration of the extrinsic coagulation pathway in Teleostei: identification of zebrafish coagulation factor VII. Proc. Natl. Acad. Sci. U.S.A. 98, 8768–8773. 10.1073/pnas.131109398.

Shimomura, T., Kondo, J., Ochiai, M., Naka, D., Miyazawa, K., Morimoto, Y., Kitamura, N., 1993. Activation of the zymogen of hepatocyte growth factor activator by thrombin. J. Biol. Chem. 268, 22927–22932. 10.1016/S0021-9258(18)41615-8.

Tsutsui, S., Yoshimura, A., Iwakuma, Y., Nakamura, O., 2023. Discovery of teleost plasma kallikrein/coagulation factor XI-like gene from channel catfish (*Ictalurus punctatus*) and the evidence that the protein encoded by it acts as a lectin. J. Mol. Evol. 91, 536–551. 10.1007/s00239-023-10113-4.

Wong, M.K.S., Takei, Y., 2013. Lack of plasma kallikrein-kinin system cascade in teleosts. PLoS One 8, e81057. 10.1371/journal.pone.0081057.

